# MEMS-Based Ultrasonic Energy Harvesting Platform Enabling Sustained *In Vivo* Operation of Implantable Microdevices

**DOI:** 10.64898/2026.07.15.738778

**Authors:** Xu Tian, Argyris Spyrou, Theocharis Nikiforos Iordanidis, Göran Stemme, Niclas Roxhed

## Abstract

Implantable microdevices capable of autonomous operation over extended lifetimes are promising enablers for minimally invasive diagnostics and therapy. Microelectromechanical systems (MEMS)-based piezoelectric ultrasonic energy harvesters (PUEH) have emerged as a compelling approach for powering implantable microdevices, where both miniaturization and efficient wireless energy transfer are essential. Here, we present a highly miniaturized (5 × 5 × 5 mm³) ultrasonic energy-harvesting platform enabling sustained *in vivo* operation of implantable microdevices. The platform integrates a MEMS-PUEH, a high-efficiency power management system, an energy storage element, and representative functional electronics. We first investigate the effect of backside cavity boundary conditions on MEMS-PUEH performance and show that a sealed air-filled chamber significantly outperforms an open water-filled cavity, yielding a 46% increase in root-mean-square output voltage and a 117% increase in average output power across a 2 kΩ resistive load under identical incident acoustic intensity at the respective optimal operating frequencies. We then demonstrate system-level integration and characterization. In a tissue-mimicking phantom, under an incident acoustic intensity of approximately 257 mW/cm^2^, the device charges an 11.5 mF supercapacitor, a 5 µAh solid-state microbattery, and a 100 µF capacitor to their nominal voltages in less than 5 min, 3 min, and 20 s, respectively. Finally, *in vivo* validation demonstrates fully autonomous operation of representative functional electronics following ultrasonic charging of the onboard energy storage element. These results establish a highly miniaturized and fully integrated ultrasonic energy-harvesting platform that advances MEMS-based power solutions for implantable biomedical microdevices.

## 1. Introduction

The rapid expansion of implantable medical devices (IMD) for extended and minimally invasive diagnostic and therapeutic applications, including implantable biosensors, neural modulators, and cardiac pacemakers, has created a growing demand for sustained, safe, and reliable power sources in miniaturized formats^[1–4]^. Advances in microelectronics^[5]^, sensor technologies^[6]^, and biointerface materials^[7]^ have enabled increasingly compact implantable platforms for diverse biomedical applications. However, the broader clinical translation of these devices remains challenging due to the lack of integrated power solutions that simultaneously provide sustained operational lifetime and small device footprints^[8]^. To overcome these limitations, wireless power transfer techniques have been widely explored as alternatives or complements to conventional batteries for IMD.

Among the wireless power transfer techniques explored for IMD, electromagnetic techniques such as inductive coupling and radiative far-field, together with acoustic approaches such as ultrasonic energy transfer, are the most widely studied^[9]^. While electromagnetic-based methods can achieve high transfer efficiencies over short distances, their performance typically deteriorates with increasing implantation depth due to stronger tissue absorption and sensitivity to positional misalignment, limiting reliability under deep or dynamic implantation conditions^[2]^. Owing to shorter wavelengths and lower ultrasound attenuation in biological tissue, ultrasonic energy transfer can support smaller receiver dimensions while maintaining effective energy delivery at clinically relevant depths, making it a strong candidate for deeply implanted and highly miniaturized devices^[10]^. Furthermore, the long-standing use of ultrasound in medical applications provides a strong foundation for safe operation under controlled acoustic intensities. These characteristics make ultrasonic energy transfer particularly well-suited for powering compact implantable microdevices through soft tissue^[11,12]^.

Ultrasonic energy transfer for implantable applications is commonly implemented using piezoelectric ultrasonic energy harvesters (PUEH), which generate electrical energy by converting incident acoustic pressure into mechanical deformation and, subsequently, electrical charge through the direct piezoelectric effect^[13]^. In recent years, a wide range of PUEH implementations for biomedical applications have been reported, including piezoelectric nanogenerators^[14–16]^, piezoelectric crystals^[17–19]^, piezoelectric composite materials^[20–22]^, and piezoelectric diaphragms^[23–32]^. As implantable systems continue to miniaturize while increasing in functionality, microelectromechanical systems (MEMS)-based PUEH have emerged as a promising solution due to their resonance tunability, precise dimensional control, batch fabrication capability, and compatibility with integrated circuits^[33,34]^. These devices typically employ thin diaphragms composed of a piezoelectric active layer bonded to a passive structural layer, such as silicon, with patterned electrodes for charge collection^[35]^. This architecture enables resonance tuning in the hundreds-of-kilohertz range, balancing efficient acoustic coupling with reduced tissue attenuation^[36]^. Importantly, wafer-level MEMS fabrication allows scalable production of highly reproducible devices with miniaturized dimensions well aligned with the constraints of minimally invasive implantation^[37]^. Notably, recent studies have shown that MEMS-PUEH can deliver power outputs on a similar scale to other reported solutions^[29,30]^ and be capable of meeting the power requirements of modern IMD^[3]^. Together, these advantages strengthen the potential of MEMS-PUEH to bridge efficient ultrasonic power transfer with the miniaturization, scalability, and system compatibility needed for future IMD.

Recent progress in MEMS-PUEH has expanded their capabilities through advances in device design, material integration, and fabrication strategy. On the device side, geometric optimization has been shown to broaden the operating bandwidth and improve harvesting performance^[25]^. MEMS diaphragm structure has been systematically investigated for low-frequency operation and resistance to misalignment and orientation variations^[36]^. In parallel, integrating bulk PZT material into MEMS-PUEH has demonstrated high energy conversion efficiency^[29]^, whereas lead-free materials such as AlN have attracted increasing interest because of their biocompatibility and CMOS compatibility^[26,27]^. At the same time, low-temperature device fabrication strategies have further enabled highly miniaturized devices with sub-milliwatt output under ultrasound intensities below regulatory safety limits^[28,30]^. Despite these progresses, the field has remained largely focused on transducer-level demonstrations. Practical implantable use, however, requires a fully integrated and highly miniaturized platform in which energy harvesting is coupled with power regulation and management, energy storage, and functional device operation^[1]^. To date, such system-level integration under physiologically relevant conditions has not been established for MEMS-PUEH, and *in vivo* validation has remained absent. Moreover, backside cavity boundary conditions represent an important yet underexplored design parameter in MEMS-PUEH, as they can critically affect damping, resonance characteristics, and harvested electrical output. Since the clinical relevance of MEMS-PUEH depends not only on device performance but also on system-level demonstration under physiological environments, advancing the field requires the development of a platform that combines a MEMS-PUEH with optimized backside cavity boundary conditions, full system integration, and *in vivo* validation for sustained functional operation of implantable microdevices.

In this work, we present a highly miniaturized MEMS-based piezoelectric ultrasonic energy harvesting platform for sustained *in vivo* operation of implantable microdevices. The platform integrates a MEMS-PUEH, power management, energy storage, and representative functional electronics within a compact 5 × 5 × 5 mm³ form factor. We investigate cavity-dependent MEMS-PUEH performance, evaluate system-level operation in a tissue-mimicking phantom, and validate autonomous operation *in vivo*. Together, these results advance MEMS-based ultrasonic energy harvesting from a transducer-level concept to a fully integrated implantable platform. More broadly, it provides a foundation for future self-contained biomedical microsystems enabled by MEMS technology, with extended operational lifetime, increased functional complexity, and improved prospects for clinical translation.

## 2. Results and Discussion

### 2.1 Integrated MEMS-based ultrasonic energy-harvesting platform

The platform is designed as a compact modular power solution for implantable microdevices, with the aim of enabling sustained autonomous device operation *in vivo* following ultrasonic energy transfer (Figure 1a). At the system level, the main design consideration is to incorporate energy harvesting, power management, energy storage, and functional electronics within a single platform while maintaining a form factor suitable for minimally invasive implantation. Realization of the complete system in a 5 × 5 × 5 mm³ form factor demonstrates the feasibility of combining multiple heterogeneous subsystems into a unified architecture with a high degree of miniaturization.

**Figure 1.**
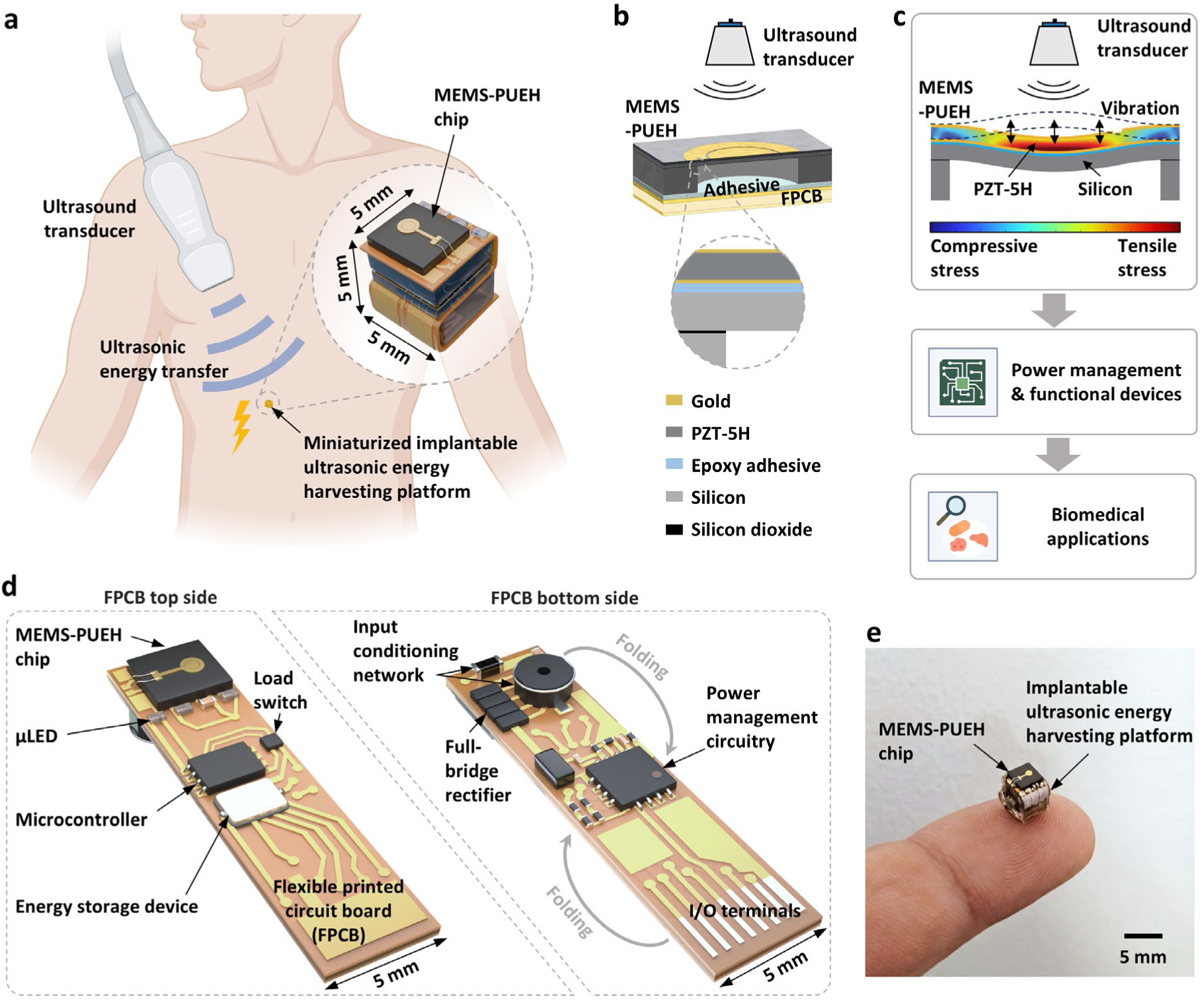
MEMS-based ultrasonic energy-harvesting platform enabling sustained operation of implantable microdevices *in vivo*. **(a)** Schematic illustration of the highly miniaturized ultrasonic energy-harvesting platform (5 × 5 × 5 mm³), which converts incident ultrasonic energy into electrical energy through a MEMS-PUEH, for biomedical applications. **(b)** Cross-sectional schematic illustration of the MEMS-PUEH structure. **(c)** Schematic illustration of the MEMS-PUEH working principle and its integration with the power management and functional electronics units for energy harvesting and delivery to biomedical microdevices. **(d)** Exploded schematic illustration of the device architecture, showing the FPCB-based integration of the MEMS-PUEH, the power management, and the functional electronics. **(e)** Photograph of the fully assembled ultrasonic energy-harvesting platform.

The energy transduction element is a MEMS-PUEH chip based on a multilayer diaphragm structure consisting of gold electrodes, a bulk PZT-5H piezoelectric layer, an adhesive layer, and a silicon structural layer (Figure 1b). Under ultrasonic excitation, the diaphragm undergoes mechanical vibration at resonance, inducing mechanical stress in the piezoelectric layer (Figure 1c). Through the direct piezoelectric effect, this stress is converted into an alternating electrical output. The generated energy is then processed by the onboard power management electronics and stored in the energy storage device, which provides sustained power for the implantable microdevice.

The overall architecture follows a modular design strategy implemented on a flexible printed circuit board (FPCB), which serves as the system-level integration platform for the energy harvesting unit, the power management unit, and the functional electronics unit (Figure 1d). The power management unit includes the input-conditioning network, full-bridge rectifier, power management circuitry, and energy storage device, whereas the functional electronics unit comprises the load switch, low-power microcontroller, and µLED. The developed platform achieves a high degree of miniaturization, modularity, and functional integration, while the folding configuration further enables efficient three-dimensional packaging without compromising electrical connectivity or mechanical stability (Figure 1e).

### 2.2 MEMS-PUEH with different backside cavity boundary conditions

To examine the effect of backside cavity boundary conditions, we compared two otherwise identical MEMS-PUEH configurations: an open water-filled cavity design^[30]^ (version 1) and a sealed air-filled chamber design (version 2), which is the configuration adopted in the integrated energy-harvesting platform (Figure 2a). This comparison isolates the influence of backside cavity boundary conditions while keeping the overall device structure unchanged. Electrical measurements were performed using the setup in Figure 2b under controlled ultrasonic excitation and defined alignment conditions.

**Figure 2.**
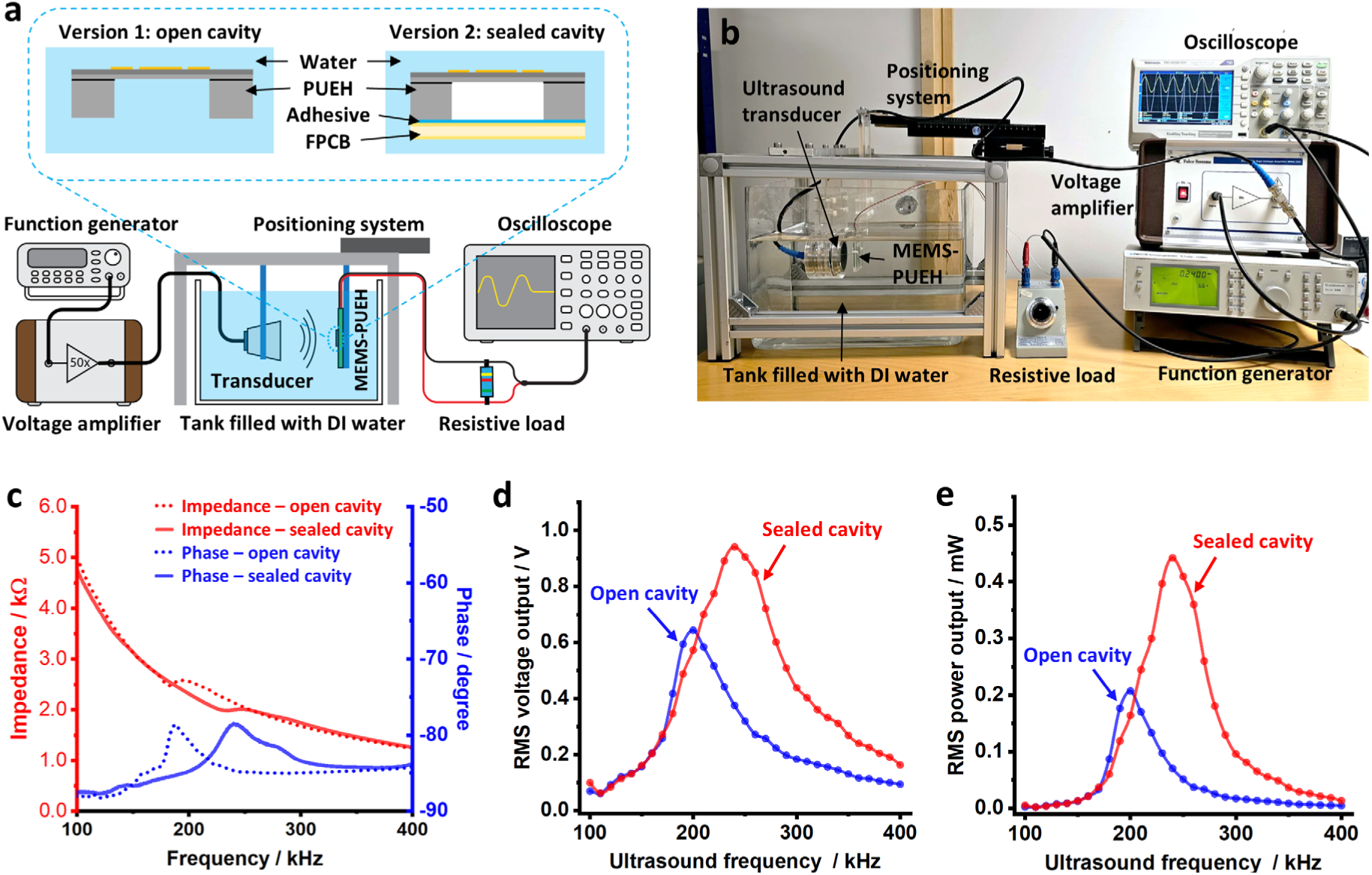
Comparative study of MEMS-PUEH with different backside cavity boundary conditions. (a) Schematic illustrations of the MEMS-based PUEH with an open water-filled cavity (version 1) and a sealed air-filled cavity (version 2), together with the associated experimental configuration used for electrical measurements. (b) Photograph of the experimental setup for electrical characterization of the MEMS-PUEH in water, showing the electrical connections used to measure voltage and power output across a resistive load. (c) Measured impedance magnitude and phase responses of the two MEMS-PUEH configurations. (d) Measured root-mean-square (RMS) output voltage across a 2 kΩ resistive load for the two MEMS-PUEH configurations, tested in water at their respective resonance frequencies of 200 kHz for the open water-filled cavity and 240 kHz for the sealed air-filled cavity under an identical incident acoustic intensity of 178 mW/cm². (e) Measured average output power across a 2 kΩ resistive load for the two MEMS-PUEH configurations under the same test conditions.

The impedance measurement results shown in Figure 2c reveal that the backside cavity condition has a pronounced influence on the mechanical resonance of the MEMS-PUEH. For the device with the open water-filled cavity, a clear resonance was observed at approximately 190–200 kHz, with an impedance magnitude of about 2.5 kΩ. This measured resonance is in good agreement with the finite-element prediction of 185 kHz for a device incorporating a 17 µm-thick PZT-5H layer^[30]^. The associated phase transition, from about −88° off resonance to around −78° near resonance, is consistent with the expected reduction in the reactive contribution as the device approaches resonant operation^[30]^. When the backside cavity was replaced by a sealed air-filled chamber, the resonance shifted upward to approximately 230– 240 kHz and the impedance decreased to about 2.0 kΩ. This shift is attributed to the reduced mass loading on the vibrating diaphragm compared with the water-filled cavity configuration. Notably, the phase response remained similar in form, indicating that the electromechanical coupling mechanism itself was preserved, while the boundary condition altered the effective dynamic loading and therefore the resonance characteristics.

The change in the resonance characteristics of MEMS-PUEH with a sealed air-filled chamber directly translated into improved electrical performance. Based on the impedance measurement results for the two backside cavity configurations, a 2.0 kΩ resistive load was selected for electrical output measurements. Under identical test conditions in water, with the ultrasound transducer and MEMS-PUEH aligned in parallel at a fixed separation distance of 20 mm and an incident acoustic intensity of 178 mW/cm² measured by a calibrated hydrophone, the two devices were evaluated at their respective resonance frequencies: 200 kHz for the open-cavity device and 240 kHz for the sealed air-filled device. Under these matched excitation conditions, the sealed air-filled chamber configuration exhibited a 46% increase in root-mean-square output voltage and a 117% increase in average output power relative to the water-filled cavity device, as shown in Figures 2d and 2e, respectively. The markedly enhanced output of the MEMS-PUEH with a sealed air-filled chamber can be attributed to the distinct mechanical environments imposed by the two backside cavity conditions. In the sealed air-filled device, the air chamber effectively isolates the backside of the diaphragm from the surrounding fluid. This reduces acoustic damping and fluid loading, allowing larger diaphragm vibration amplitude and a more pronounced resonant response, which in turn enhances electromechanical energy conversion. In contrast, when the backside cavity is filled with water, the diaphragm is subjected to additional fluid loading that limits its motion and reduces the electrical output of the device. These results therefore show that the air-filled backside cavity is not merely a packaging solution, but a key design parameter governing both resonance tuning and electrical performance of the MEMS-PUEH.

### 2.3 System-level integration of the ultrasonic energy harvesting platform

The progression from a high-performance MEMS-PUEH to a functional implantable energy-harvesting platform capable of supporting sustained *in vivo* operation of microdevices is presented in Figure 3. The advancement at this stage lies in integrating the MEMS-PUEH chip with a system architecture that manages energy in a form usable by implantable microdevices (Figure 3a). The power management unit conditions, regulates, and stores the harvested electrical energy. Because the MEMS-PUEH behaves as a high-impedance and predominantly capacitive source near its resonance frequency, an input conditioning network is placed directly at its output. This network is designed to compensate for the capacitive behavior of the MEMS-PUEH and promote voltage buildup at the interface between the MEMS-PUEH and the following full-bridge rectifier. This voltage-boosting function is particularly important for maintaining a sufficient level of input voltage, thereby ensuring a reliable cold start of the power management integrated circuit (PMIC) under variable acoustic input intensities and coupling conditions during ultrasonic energy transfer.

**Figure 3.**
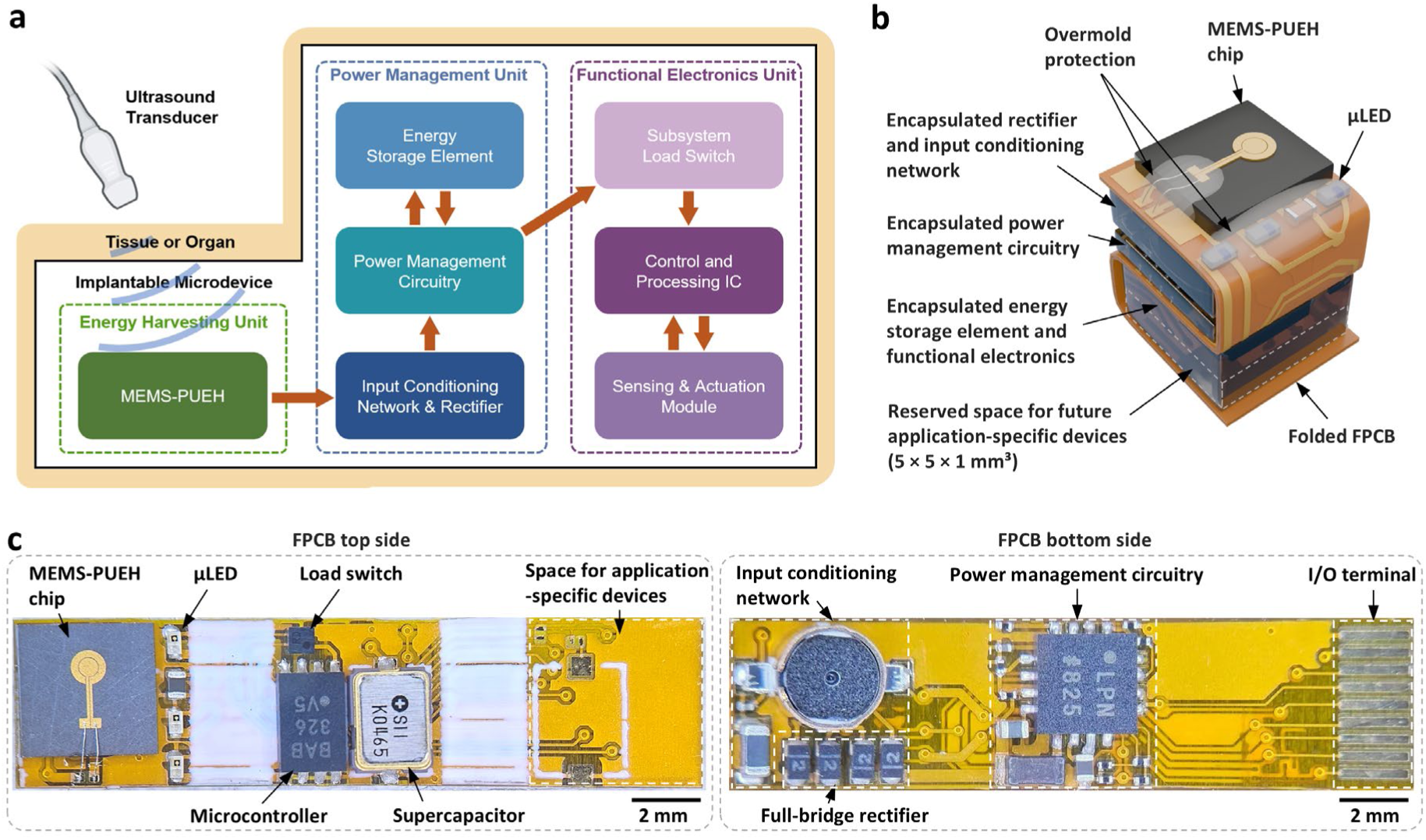
System-level integration and assembly of the MEMS-based ultrasonic energy harvesting platform. (a) System block diagram illustrating the energy conversion, management, storage, and load-delivery pathway for autonomous operation of the implantable microdevice. (b) Schematic illustration of the assembled platform, showing the three-dimensional layout and modular integration of the device components. (c) Photographs of the top and bottom sides of the assembled FPCB.

The conditioned alternating electrical output is then converted into a unidirectional input voltage by a full-bridge rectifier. Full-wave rectification ensures that energy from both the positive and negative half-cycles of the MEMS-PUEH output contributes to the system input, which is particularly beneficial under low-input-energy conditions. The rectified output is supplied to the power management circuitry, which regulates the rectified voltage and coordinates energy transfer among the MEMS-PUEH, the energy storage element, and the subsequent functional electronics unit. The harvested energy is first stored in an onboard energy storage element, which decouples the continuous but low-power energy-harvesting process from the intermittent and higher peak-power demands of the electronic load. As the energy storage element voltage increases, the PMIC monitors predefined voltage thresholds to determine when sufficient energy has been accumulated for a stable device operation. Once the energy storage element is charged to operational terminal voltage, the PMIC enables the functional electronics unit, triggering a light indication from the electronic subsystem indicating that the device is ready for autonomous *in vivo* operation.

The functional electronics unit acts as a representative demonstration of the use of the ultrasonic energy harvested and stored by the platform. A load switch is incorporated to provide controlled power delivery to a microcontroller-driven functional subsystem. By isolating the load during the energy accumulation phase, the load switch prevents premature discharge of the energy storage element and ensures that the functional subsystem is activated only when adequate energy is available. Once the load switch is enabled by PMIC, the operation of the functional subsystem is initiated. The functional subsystem consists of a low-power microcontroller and an array of µLEDs, representing implantable electronic systems capable of controlling, processing, and signaling. Upon activation, the microcontroller executes its programmed task, while the µLED provides a clear indication of successful ultrasonic energy harvesting, energy storage element charging, and energy delivery to the load. As the stored energy falls below the predefined threshold, the functional subsystem is disconnected to prevent further depletion of the energy storage element, and the element can be recharged again through ultrasonic energy transfer, enabling long-term cyclic operation of the implantable microdevice.

Figure 3b further presents a schematic of the fully assembled system architecture, highlighting the modular integration enabled by the FPCB and demonstrating the packaging solutions implemented for implantable applications. The electronic components are encapsulated with medical-grade epoxy for electrical insulation and environmental protection, while the protruding wire bonds on the MEMS-PUEH and µLEDs are mechanically protected by overmolding. The entire device was then coated with a parylene-C layer for biocompatibility. Beyond the representative functional electronics demonstrated, the platform can accommodate future application-specific modules for target biomedical applications through the reserved space (approximately 5 × 5 × 1 mm³). Figure 3c shows photographs of the top and bottom sides of the FPCB with all the system components installed, demonstrating the successful integration of the MEMS-PUEH unit, power management unit, and functional electronics unit within a single compact platform.

### 2.4 System-level parametric characterization of ultrasonic energy harvesting

To evaluate the system-level energy transfer performance of the MEMS-based ultrasonic energy harvesting platform, we characterized the voltage and power delivered to the power management circuitry under controlled acoustic conditions. For this study, a MEMS-PUEH with an air-filled chamber was immersed in a water tank and electrically connected to the remaining platform components (Figure 4a). This arrangement enabled direct measurement of incident acoustic intensity, precise adjustment of MEMS-PUEH orientation, and accurate control of the transducer-to-harvester separation distance.

**Figure 4.**
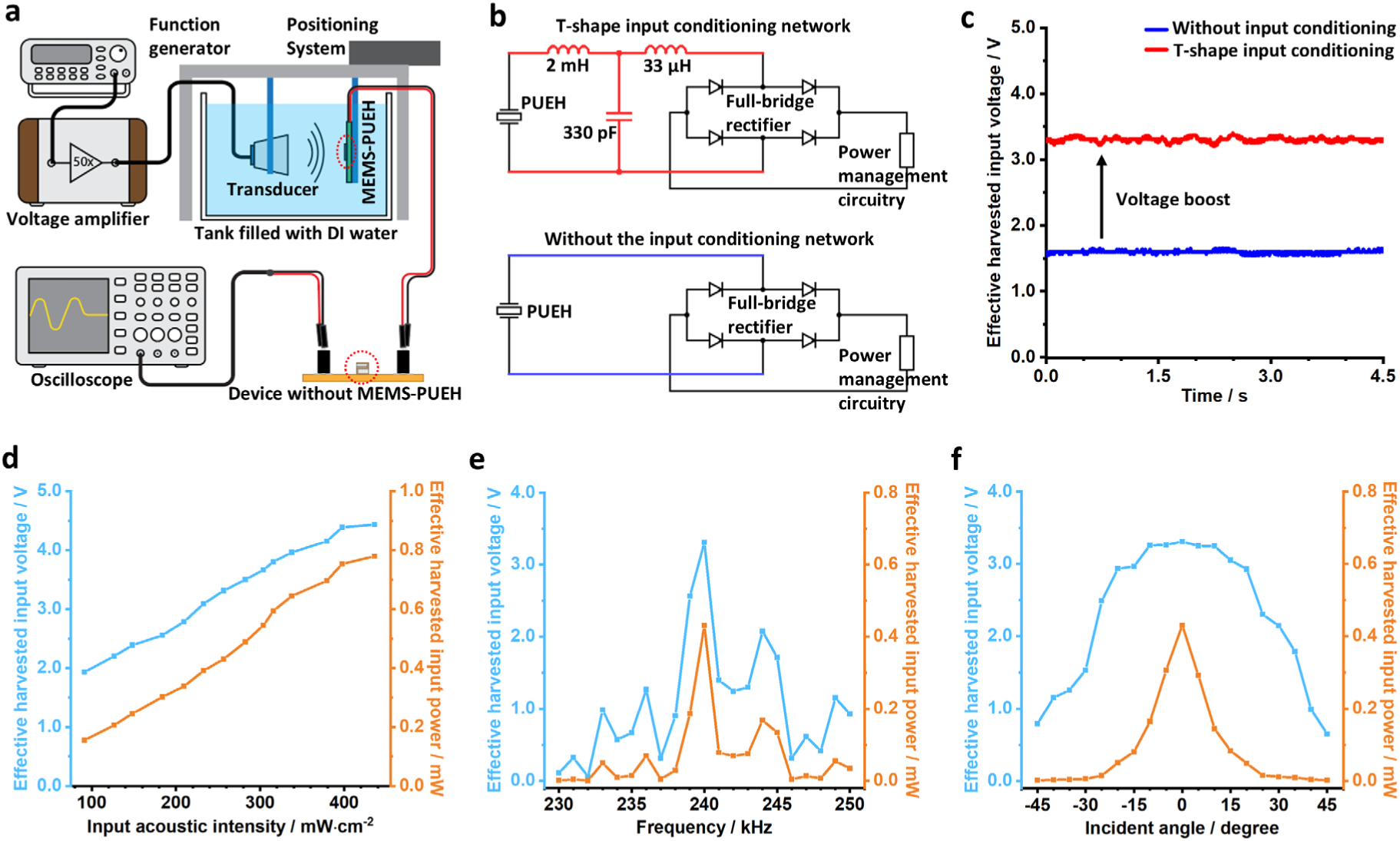
System-level parametric characterization of MEMS-based ultrasonic energy harvesting platform: **(a)** Schematic illustration of the experimental setup used for system-level parametric characterization. **(b)** Circuit diagram of the MEMS-PUEH interface with and without a T-shaped input conditioning network, followed by a full-bridge rectifier and the power management circuitry represented as an equivalent load. **(c)** Measured effective harvested input voltage with and without input conditioning, indicating improved electrical coupling and a more reliable cold start of the system under *in vivo* operating conditions. (d) Measured effective harvested input voltage and effective harvested input power as functions of incident acoustic intensity at the resonance frequency of 240 kHz, with the MEMS-PUEH aligned parallel to the ultrasound transducer. (e) Measured effective harvested input voltage and effective harvested input power as functions of ultrasound frequency at a fixed incident acoustic intensity of 257 mW/cm², with the MEMS-PUEH aligned parallel to the ultrasound transducer. (f) Measured effective harvested input voltage and effective harvested input power as functions of incident angle at the resonance frequency of 240 kHz and a fixed incident acoustic intensity of 257 mW/cm².

The MEMS-PUEH was connected to a full-bridge rectifier and the following power management circuitry, either directly or through a T-shaped input conditioning network (Figure 4b). The electrical quantities measured at the input of the power management circuitry were used here as system-level performance metrics because they represent the usable electrical output available to the PMIC and therefore directly determine its operation and the subsequent charging of the energy storage element. Incorporating the T-shaped conditioning network increased the effective harvested input voltage from 1.6 V to 3.3 V under an identical incident acoustic intensity of 257 mW/cm² at the MEMS-PUEH surface, corresponding to a 106% improvement (Figure 4c). This substantial increase indicates that the network promotes voltage buildup at the rectifier input and provides a more favorable operating condition for the PMIC. In practical terms, this makes the conditioning network an important enabling element for reliable cold start and stable energy transfer under variable *in vivo* operating conditions.

The amount of usable electrical input delivered to the PMIC also scaled strongly with the applied acoustic excitation. At 240 kHz, both harvested input voltage and harvested input power increased with incident acoustic intensity over the investigated range of 91 mW/cm² to 436 mW/cm². The harvested input voltage increased from 1.9 V to 4.4 V, while the harvested input power increased from 0.15 mW to 0.78 mW (Figure 4d). This trend indicates that, within this operating window, increasing acoustic drive effectively translates into greater electrical input to the power management circuitry. At the same time, optimum operation required more than increased acoustic intensity alone. We evaluated the frequency dependence of energy transfer to the power management stage under a fixed incident acoustic intensity of 257 mW/cm². This intensity was selected because it produced an input voltage of 3.3 V at the resonance frequency of 240 kHz, corresponding to the upper limit of the recommended input range of the PMIC. As presented in Figure 4e, both the effective harvested input voltage and the effective harvested input power exhibited a pronounced frequency-dependent response, with maximum values occurring near the mechanical resonance of the MEMS-PUEH at 240 kHz. At resonance, the harvested input voltage reached 3.3 V, and the harvested input power reached 0.43 mW. These results indicate that efficient energy delivery to the platform is strongly dependent on proper frequency tuning. Angular alignment introduced a further practical constraint, although the platform showed useful tolerance to moderate misalignment (Figure 4f). The harvested input voltage remained close to 3.3 V for angular misalignment below 20°, while the harvested input power decreased gradually as the angle increased from 0° to 30°. Once the misalignment became large, however, the degradation was severe: at 45°, the harvested input voltage dropped to 0.70 V and the harvested input power fell below 2 µW. These results indicate that small angular deviations do not immediately compromise PMIC input conditions, whereas larger angular offsets sharply reduce acoustic coupling and therefore limit the electrical energy delivered to the downstream circuitry.

Taken together, these results show that the system-level performance of the platform is governed by a coupled interaction between the electrical interface and the acoustic operating conditions. The T-shaped conditioning network makes the harvested energy substantially more accessible to the PMIC, while incident acoustic intensity, excitation frequency, and angular alignment determine how efficiently acoustic energy is converted into usable electrical input under a given condition. For the present platform, the most favorable operating condition is obtained near 240 kHz with an incident acoustic intensity of 257 mW/cm² and minimal angular misalignment (< 20°), where efficient PMIC input delivery is achieved without exceeding the recommended input voltage range.

### 2.5 System-level demonstration in a tissue-like phantom model and *in vivo*

To demonstrate system-level functionality of the MEMS-based ultrasonic energy harvesting platform, we performed experiments in a tissue-like gelatin-based phantom model and *in vivo*. These studies were designed to evaluate the performance of the platform in charging the integrated energy storage elements via ultrasonic energy transfer and to verify that the fully integrated system can harvest, store, and deliver energy to the functional electronics under physiologically relevant conditions.

Figure 5a presents a schematic illustration of the experimental setup used for phantom-based characterization. A positioning system was implemented to enable precise adjustment of the relative position between the ultrasound transducer and the energy harvesting platform. Figure 5b shows a photograph of the device embedded in the tissue-like gelatin-based phantom model, where the illuminated µLED indicates successful system-level operation powered solely by the harvested and stored ultrasonic energy. The separation distance between the ultrasound transducer and the device was fixed at 20 mm. At the same distance in water, the incident acoustic intensity at 240 kHz frequency was measured to be 257 mW/cm². In the gelatin-based phantom model, the incident acoustic intensity is expected to be slightly lower due to the higher acoustic attenuation of the medium. To demonstrate the versatility of the platform, three different energy storage elements were integrated and evaluated in the phantom, including a 100 µF ceramic capacitor, a 5 µAh solid-state microbattery, and an 11.5 mF supercapacitor (Figure 3c). During ultrasonic charging, the terminal voltage of each integrated energy storage element was monitored over time, and the corresponding charging profiles are shown in Figure 5c. All three devices were successfully charged to their respective nominal voltages via ultrasonic energy transfer, although the charging profiles differed substantially because of their different storage capacities and nominal voltages. The 100 µF ceramic capacitor was charged to its nominal voltage of 4.0 V within 16 s, while the 5 µAh solid-state microbattery reached its nominal voltage of 3.8 V after 2 min 8 s, and lastly, the 11.5 mF supercapacitor showed a longer charging duration, reaching its nominal voltage of 2.6 V after 4 min 41 s. These results demonstrate that the presented energy-harvesting platform is compatible with a broad range of energy storage elements, supporting application-specific configuration of the overall implantable microdevice.

**Figure 5.**
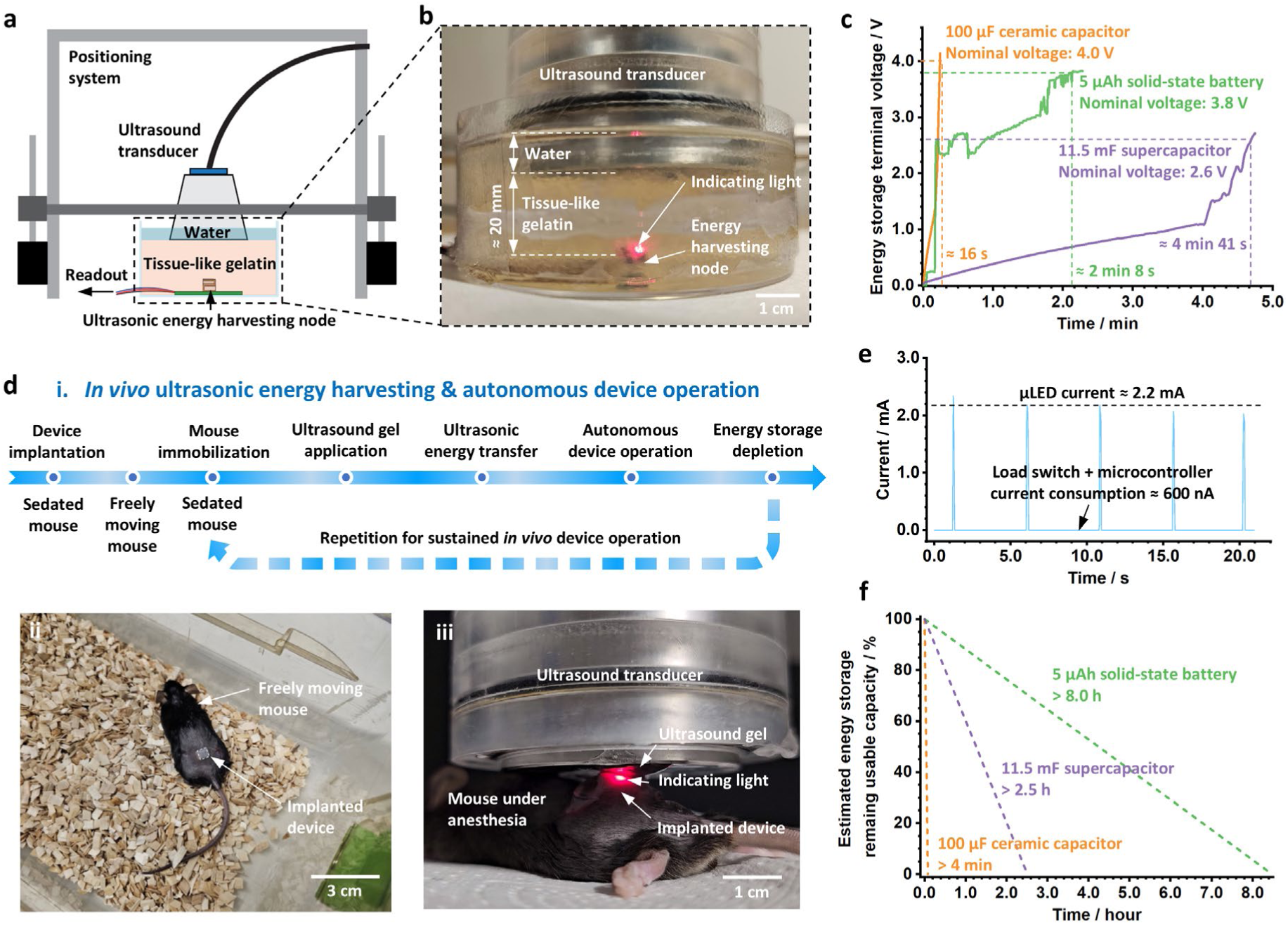
System-level demonstration of the MEMS-based ultrasonic energy harvesting platform in a tissue-like gelatin phantom model and *in vivo*. (a) Schematic illustration of the experimental setup used for validation in the gelatin-based tissue-like phantom model. (b) Photograph of the phantom-model setup used for electrical characterization of the energy harvesting platform. (c) Measured terminal voltage as a function of charging time for three integrated energy storage elements in the gelatin-based phantom model: a 100 µF ceramic capacitor (4.0 V nominal voltage), a 5 µAh solid-state microbattery (3.8 V nominal voltage), and an 11.5 mF supercapacitor (2.6 V nominal voltage). (d) *In vivo* validation of the developed platform: (i) Experimental scheme of *in vivo* ultrasonic energy charging of the energy storage element, followed by autonomous device operation. (ii) A freely moving mouse carrying a subcutaneously implanted microdevice. (iii) Successful ultrasonic charging of a 100 µF ceramic capacitor and autonomous operation of the implanted microdevice, as verified by µLED blinking. (e) Measured current consumption of the onboard functional electronics powered by a fully charged 11.5 mF supercapacitor at its nominal voltage of 2.6 V, showing that the average quiescent current of the load switch and the microcontroller in low-power mode is approximately 600 nA. (f) Calculated theoretical *in vivo* operating time of the microsystem with the representative functional in idle states when powered by different fully charged energy storage elements.

To further assess system-level functionality under physiological conditions, the device was evaluated *in vivo* following subcutaneous implantation (Figure 5d). Figure 5d-i illustrates the sequence of the experimental procedure for device implantation and *in vivo* ultrasonic energy harvesting. Following depletion of the stored energy, the ultrasonic charging process can be repeated to restore device operation, enabling on-demand ultrasonic energy transfer for sustained *in vivo* operation. Because of the short charging time, such charging could in principle be performed with minimal intervention, for example, by brief immobilization of the animal during ultrasonic charging through physical restraint without anesthesia^[38]^. As illustrated in Figure 5d-ii, the small overall device form factor underscores its applicability for minimally invasive implantation and prospective operation in freely moving animal models^[39]^. As shown in Figure 5d-iii, ultrasonic energy transfer successfully charged a 100 µF ceramic capacitor integrated in the device and enabled autonomous operation of the implanted microdevice, as verified by µLED blinking. The incident acoustic intensity is estimated to be approximately 400 mW/cm² based on measurements at the same distance in water, well below the FDA safety limit of 720 mW/cm². Although these charging conditions may transiently cause an overshoot beyond the nominal maximum input voltage of the PMIC, no device failure was observed, likely due to the short charging duration (≈ 15 s) and the low input power (< 0.8 mW). Further reductions in charging time are expected through optimization of the ultrasound source and improved transmitter-device alignment. These results provide a system-level validation that the developed integrated ultrasonic harvesting platform can harvest, store, and deliver energy under physiological conditions, supporting its potential as a wireless power solution for compact implantable microdevices.

We further characterized the power consumption of the functional electronics unit, which serves as a representative system load. Figure 5e shows the measured current profile of the functional electronics unit powered by the fully charged 2.6 V supercapacitor in the phantom model. With the load switch enabled and the microcontroller in low-power mode, the functional electronics unit exhibited an average standby current of approximately 600 nA. Upon task execution, the current increased to 2.2 mA due to microcontroller operation and µLED illumination. These measurements define the power consumption profile of a typical functional load and provide the basis for estimating the theoretical *in vivo* operating time of the implantable device after a complete ultrasonic energy transfer cycle. Including the 260 nA quiescent current of the PMIC, the total standby current of the platform was approximately 860 nA, corresponding to a power consumption of about 2.24 µW at 2.6 V at the standby state (energy storage element is charged at operational terminal voltage, load switch is on, and microcontroller is in low-power mode with watchdog timer on). Using the 2.24 µW power consumption and the 1.8 V minimum operating voltage of the microcontroller as the estimation framework, the theoretical operating time frames supported by the different energy storage devices were calculated, as presented in Figure 5f. For a fully charged 100 µF capacitor and a fully charged 11.5 mF supercapacitor, the operating times were estimated from their usable voltage windows and exceeded 4 min and 2.5 h, respectively. For a fully charged 5 µAh solid-state microbattery, the operating time was estimated from its nominal capacity and exceeded 8 h. For System-on-Chips (SoCs) operating at ultra-low power levels of 1 µW or below^[40]^, the supported operating time would be extended further. In this regime, the 5 µAh solid-state microbattery could sustain operation for about 19 h, and even longer durations approaching or exceeding one day could be expected for sub-µW loads. Together with its cyclic recharging capability, these position the MEMS-based platform as a valuable power solution for long-term implantable microdevices.

## 3. Conclusions

In conclusion, we developed a highly miniaturized MEMS-based ultrasonic energy harvesting platform for sustained operation of implantable microdevices. The system integrates a MEMS-PUEH, power management system, energy storage element, and representative functional electronics within a compact 5 × 5 × 5 mm³ form factor, thereby extending MEMS-based PUEH from a transducer-level demonstration to an implantable microsystem with validated functionality. We show that cavity boundary conditions strongly influence harvester performance, with the sealed air-filled chamber configuration yielding substantially higher voltage and power output than the open water-filled cavity. The integrated platform further demonstrated efficient charging of multiple energy storage elements in a tissue-mimicking phantom and enabled fully autonomous operation of a microcontroller-driven representative implant load *in vivo* following ultrasonic charging. These results demonstrate that the MEMS-based ultrasonic energy harvesting platform is capable of enabling the sustained, functional operation of implantable microdevices under physiologically relevant conditions. Overall, our work establishes a foundation for MEMS technology-enabled self-contained biomedical microsystems with extended operational lifetime, increased functional complexity, and improved prospects for clinical translation.

## Materials and Methods

### Fabrication of the MEMS-PUEH chip

The MEMS-PUEH chip was fabricated as described earlier^[30]^. In short, a bulk PZT-5H sheet (T105-H4NO-2929, Mide Technology, USA) was polished and metallized with a TiW/Au (50 nm/600 nm) for creating the floating electrode of the MEMS-PUEH. An epoxy adhesive layer (ECO-System, Epodex, Germany, mixing ratio 2:1) was spin-coated onto an SOI wafer (22 µm device Si / 1 µm BOX / 300 µm handle). The metalized side of the bulk PZT-5H sheet and the SOI wafer were then bonded together using low temperature and pressure (80 °C, 1.5 MPa) for 2 h. The PZT-5H was manually thinned and polished to a thickness of approximately 17 µm. A cavity (900 µm diameter) on the backside was created using SiO_2_ as a hard mask, followed by DRIE to the BOX layer and oxide removal. Front-side electrodes (TiW/Au, 50 nm/600 nm) were patterned via a lift-off process. All processing temperatures were kept below 85 °C. The wafer stack was diced into individual chips measuring 4 × 4 mm^2^ for subsequent electrical characterizations and device integration.

### Impedance measurement of the MEMS-PUEH chip

The diced MEMS-PUEH chip was mounted onto a customized printed circuit board (PCB) (FR4, JDB Technology, China) using glue (Rubberfix, Casco, Sweden), and electrical interconnections were established via wire bonding (4523D, Kulicke & Soffa, Singapore). The assembled device was subsequently coated with an approximately 8 μm-thick layer of parylene-C (Galentis S.r.l., Italy) using a parylene coater (PDS 2010 Labcoter 2, Parylene Deposition Systems, USA) to provide electrical insulation. The PCB was designed with a 2 mm-wide slit positioned directly beneath the MEMS-PUEH chip, enabling liquid access to the backside cavity (0.9 mm in diameter) of the device. For impedance characterization, the device was immersed in a glass water tank (30 × 22 × 24 cm³) filled with deionized (DI) water to emulate an acoustically relevant medium. The impedance characteristics of the MEMS-PUEH were measured using a frequency response analyzer (Moku:Go, Liquid Instruments, Australia). Two configurations were investigated: (i) a water-filled backside cavity and (ii) a sealed air-filled chamber. For the air-filled configuration, the backside cavity of the same device was sealed by applying a double-sided adhesive tape (Scotch, 3M, USA) together with an FPCB (Polyimide, JDB Technology, China) layer through the PCB slit, thereby preventing water ingress and maintaining an enclosed air cavity.

### Electrical characterizations of the MEMS-PUEH chip

To evaluate the electrical performance of MEMS-PUEH chips with different backside cavity boundary conditions, a piezoelectric ultrasound transducer (PA1272, 500 kHz, 44 mm, Precision Acoustics, UK) was used as the acoustic source. A 5 V peak-to-peak sinusoidal waveform in 5-cycle burst mode was generated using a function generator (PM 5139, Philips/Fluke, The Netherlands), amplified (×50) by a voltage amplifier (WMA-300, Falco Systems, The Netherlands), and applied to drive the ultrasound transducer. The ultrasound transducer and the PCB-mounted MEMS-PUEH were placed in the water tank, fixed at a separation distance of 20 mm, and aligned parallel to each other using a positioning stage. The output of the MEMS-PUEH was connected to a 2 kΩ resistive load, and the voltage across the load was recorded using an oscilloscope (TBS 1052B-EDU, Tektronix, USA). The MEMS-PUEH were tested at their respective resonance frequencies, 200 kHz for the open water cavity version and 240 kHz for the sealed air-filled chamber version, under an identical incident acoustic intensity of 178 mW/cm². The incident acoustic power intensity at the MEMS-PUEH position was measured using a needle-type hydrophone (NH1000, 1 mm, Precision Acoustics, UK) at the location of the MEMS-PUEH PZT-5H surface.

### Design and assembly of the MEMS-based ultrasonic energy harvesting platform

The assembly process was carried out on a prototype PCB breadboard. A custom-designed 100 μm-thick FPCB (polyimide, JDB Technology, China) with dimensions of 19 × 5 mm² was used as the platform for system-level integration. The FPCB was fixed onto the prototype PCB breadboard using double-sided adhesive tape and wire-bonded (4523D, Kulicke & Soffa, Singapore) to external copper tracks to enable microcontroller programming and system-level validation.

#### Energy harvesting unit

The fabricated MEMS-PUEH chip was attached to the FPCB using double-sided adhesive tape (Scotch, 3M, USA). The edges of the chip were further anchored to the FPCB and sealed using an electronics-grade silicone adhesive sealant (EGS10W-20G, Circuit Works, USA) to ensure mechanical stability and liquid resistance. Electrical connections between the top electrodes of the MEMS-PUEH and the FPCB were established via wire bonding (4523D, Kulicke & Soffa, Singapore). The bond wires were subsequently encapsulated using epoxy (301, EPO-TEK, USA) and cured at room temperature to provide mechanical reinforcement and insulation.

#### Power management unit

The design rationale of the T-shaped input conditioning network consists of two inductors (2 mH: ELT-3KN136B, Panasonic; 33 µH: MLF1608C330KTD00, TDK) and one capacitor (330 pF: CC0100KRX7R7BB331, YAGEO), with electrical connections formed using conductive epoxy (CW2460, Circuit Works, USA). A full-bridge rectifier composed of four Schottky diodes (CFSH05-20L, Central Semiconductor, USA) was implemented where electrical connections were formed using conductive epoxy. The complete power management circuit is centered around an ultralow-power boost regulator with maximum power point tracking (MPPT) and charge management (ADP5090, Analog Devices, USA), in an LFCSP_WQ package (3.00 × 3.00 × 0.75 mm³). A 11.5 mF supercapacitor (CPM3225A, Seiko Instruments, Japan) with a dimension of 3.2 × 2.5 × 0.9 mm³ was selected as a representative energy storage device. All electrical connections between the power management IC and associated passive components were realized using the conductive epoxy.

#### Functional electronics unit

An ultra-low leakage load switch, TPS22916 (Texas Instruments, USA), in a YFP0004 package (0.78 × 0.78 × 0.55 mm³), was used for controlled power delivery to the load. A low-power microcontroller (PIC12LF1522, Microchip, USA), in a UDFN package (2.00 × 3.00 × 0.50 mm³), was employed for system control and timing operations. For demonstration purposes, three red-emitting SMD LEDs (XZMDR155W, SunLED, USA), each with dimensions of 0.65 × 0.35 × 0.20 mm³, were used as representative devices. All electrical interconnections within the functional electronics unit were also established using the conductive epoxy.

#### Protection and folding

For mechanical protection and encapsulation, a 3D-printed frame (5 × 5 mm² footprint) was placed over the power management unit and functional electronics unit. The cavities were filled with encapsulant (301, EPO-TEK, USA) and cured at room temperature for 24 h. Subsequently, the FPCB was folded into a compact configuration and secured using instant glue. The fully assembled device was finally coated with an approximately 8 μm-thick layer of parylene-C to provide biocompatibility and environmental protection.

### Programming and operation of the microcontroller

The microcontroller (PIC12LF1552, Microchip, USA), an 8-pin Flash-based 8-bit CMOS microcontroller with eXtreme Low Power technology, was programmed using MPLAB X IDE v6.00 and an MPLAB PICkit 4 debugger (Microchip, USA). To minimize power consumption, the 31 kHz low-power internal oscillator was selected as the system clock. During operation, the microcontroller was configured to enter low-power sleep mode (sleep current down to 20 nA at 1.8 V, the watchdog timer current consumption down to 200 nA at 1.8 V) between active events. Periodic wake-up was triggered by the internal watchdog timer, which provides programmable time intervals from 1 ms to 256 s. After each wake-up event, the microcontroller briefly activated the LED for 10 ms to indicate that sufficient stored energy was available and to demonstrate autonomous device operation. The microcontroller then returned to sleep mode until the next watchdog-timer wake-up event. This duty-cycled operation minimized average power consumption while providing periodic visual confirmation of successful energy harvesting, storage, and controlled load activation.

### System-level characterization of the ultrasonic energy harvesting platform

To evaluate the system-level performance, a hybrid test configuration was implemented. The MEMS-PUEH was immersed in water using the positioning stage, while the remaining parts of the ultrasonic energy harvesting platform were fully assembled and placed on the prototype PCB breadboard. The device was wire-bonded to copper tracks of the prototype PCB breadboard and connected to the MEMS-PUEH output and the external acquisition system through electrical connectors. This configuration enabled accurate characterization of the acoustic input conditions (various incident acoustic intensities, frequencies, and angles) while providing direct access to the electrical nodes of the power management circuit. A piezoelectric ultrasound transducer (PA1272, 500 kHz, 44 mm, Precision Acoustics, UK) was used as the acoustic source. Peak-to-peak sinusoidal waveforms were generated using a function generator (PM 5139, Philips/Fluke, The Netherlands) at different voltage levels and frequencies to vary the incident acoustic intensity and frequency. The signals were amplified using a voltage amplifier (WMA-300, Falco Systems, The Netherlands) and applied to drive the ultrasound transducer. The ultrasound transducer and PCB-mounted MEMS-PUEH were placed in a water tank, fixed at a separation distance of 20 mm, and aligned using a positioning system. The positioning system features a rotatory nob at the beam that holds MEMS-PUEH, allowing precise adjustment of the alignment angle between the ultrasonic transducer and the MEMS-PUEH. The incident acoustic intensity was calibrated using a hydrophone (D/300, Neptune Sonar, UK). The effective input voltage to the power management IC was measured using an oscilloscope (TBS 1052B-EDU, Tektronix, USA). The corresponding input current was measured across a 10 Ω current-sensing resistor connected in series with the power management input using a multimeter (2110, Keithley, USA). The effective input power delivered to the power management circuitry was then calculated from the measured voltage and current.

### Gelatin-based phantom measurements

A positioning system consisting of an aluminum frame and a linear motion controller was used to enable precise relative positioning between the ultrasound transducer and the phantom model. The fully assembled MEMS-based ultrasonic energy harvesting platform was placed on a prototype PCB breadboard and wire-bonded to copper tracks. The opposite ends of the copper traces were soldered to external wires for real-time monitoring of the terminal voltages. The prototype PCB breadboard and device assembly were placed in a Petri dish with an extended wall formed using wrapped plastic film. The complete assembly was coated with an 8 μm-thick parylene-C layer for electrical insulation and protection. A 5% (w/v) gelatin solution was prepared to mimic soft tissue. Gelatin powder (EMPROVE^®^ ESSENTIAL, Ph. Eur., BP, NF; Merck, Cat. no. 1.04078, Stockholm, Sweden) was dissolved in deionized water and allowed to equilibrate to room temperature. The resulting gelatin precursor was then carefully poured into a Petri dish, ensuring a controlled 20 mm separation distance between the top surface of the gelatin phantom and the MEMS-PUEH chip. The phantom was subsequently left to solidify in a refrigerator at 4 °C. During experiments, a thin layer of water was added onto the solidified gelatin surface to serve as an acoustic coupling medium between the ultrasound transducer and the phantom. The ultrasound transducer was carefully positioned on the gelatin surface. A peak-to-peak sinusoidal waveform was generated using the function generator, amplified using the voltage amplifier, and applied to drive the ultrasound transducer. For reference, at the same 20 mm separation distance in water, the incident acoustic intensity at 240 kHz was measured to be 257 mW/cm² using a calibrated hydrophone (D/300, Neptune Sonar, UK). In the gelatin phantom, the incident acoustic intensity was expected to be slightly lower due to the higher acoustic attenuation of the phantom medium compared with water. The energy harvesting performance was characterized by monitoring the voltage at the BAT pin of the ADP5090 power management IC in real time during the ultrasonic energy transferring process. The 100 μF ceramic capacitor (GRM21BR60G107ME11L, Murata Electronics, Japan) and the 5 μAh solid-state microbattery (CBC005, Cymbet Corporation, USA) were monitored using a high-resolution multimeter (2110, Keithley, USA), with data recorded via KI-Tool software (Keithley Instruments, USA). The charging profile of the 11.5 mF supercapacitor (CPM3225A, Seiko Instruments, Japan) was measured using a multimeter (UT61E+, UNI-T, China), with data logged through the manufacturer’s USB communication software. To characterize the power consumption of the onboard functional electronics, the current was measured across a 10 Ω current-sensing resistor connected in series with the load switch input using the Keithley 2110 multimeter. Simultaneously, the corresponding voltage was recorded using the UNI-T multimeter. This combined measurement enabled the accurate determination of the instantaneous power consumption of the functional electronics during operation.

### *In vivo* ultrasonic energy harvesting experiments

Animal experiments were conducted in compliance with the ARRIVE guidelines and in accordance with EU Directive 2010/63/EU governing the protection and use of laboratory animals. All procedures were performed at the Comparative Medicine KM facilities at Karolinska Institutet and adhered to Swedish national regulations. Experimental protocols were reviewed and approved by the Stockholm Animal Ethics Committee under permit number 6995-2021, with amendment 00071-2024, and authorized by the Swedish Board of Agriculture. Female C57BL/6J mice aged 16–20 weeks were sourced from Charles River Laboratories (Germany). A positioning system consisting of an aluminum frame and a linear motion controller was used to enable precise relative positioning between the ultrasound transducer and the animal model. The mouse was anesthetized with isoflurane, and the flanks were shaved. A small incision was made in the right flank of each mouse to allow subcutaneous implantation of the device. The device was carefully placed into a subcutaneous pocket to ensure stable positioning beneath the skin. The skin was then closed with sutures, and the animal was transferred to a warmed recovery cage and monitored until fully recovered from anesthesia and showing no signs of distress. After recovery, the animal was re-anesthetized with isoflurane for ultrasound application. Chemolan contact gel (FS362, Chemodis, Alkmaar, The Netherlands) was then gently applied over the area of the implanted device and the surrounding skin surface to ensure adequate acoustic coupling. The ultrasound transducer was then carefully positioned above the device in contact with the ultrasound gel, with an approximate separation distance of 3 mm between the transducer surface and the skin surface. A peak-to-peak sinusoidal waveform was generated using the function generator, amplified using the voltage amplifier, and applied to drive the ultrasound transducer. For the *in vivo* experiment, the function generator output voltage was intentionally increased to improve the reliability of ultrasonic energy transfer under physiologically variable conditions, including tissue heterogeneity, device placement uncertainty, and acoustic coupling variability. For reference, at the same 3 mm separation distance in water, the incident acoustic intensity at 240 kHz was measured to be 400 mW/cm² using a calibrated hydrophone (D/300, Neptune Sonar, UK).

## Acknowledgments

This work was supported by the Swedish Foundation for Strategic Research (SSF Grant Project No. RMX18-0066). The authors would like to thank the SSF project partners and their team members for close collaboration and valuable discussions, including Prof. Per-Olof Berggren, Dr. Martin Köhler (Karolinska Institutet, Sweden), Prof. Wouter Metsola van der Wijngaart, Prof. Anna Herland (KTH Royal Institute of Technology, Sweden), and Prof. Atila Alvandpour (Linköping University, Sweden). The authors thank Prof. Elisabet Andersson and the staff of the Comparative Medicine Biomedicum (KM-B) and KM-F animal facilities for their invaluable assistance with animal care and husbandry, particularly during the mouse experiments.

